# Movement responses to environment: fast inference of variation among southern elephant seals with a mixed effects model

**DOI:** 10.1101/314690

**Authors:** I. D. Jonsen, C. R. McMahon, T. A. Patterson, M. Auger-Méthé, R. Harcourt, M. A. Hindell, S. Bestley

## Abstract

Like many species, movement patterns of southern elephant seals (*Mirounga leonina*) are being influenced by long-term environmental change. These seals migrate up to 4000 km from their breeding colonies, foraging for months in a variety of Southern Ocean habitats. Understanding how movement patterns vary with environmental features and how these relationships differ among individuals employing different foraging strategies can provide insight into foraging performance at a population level. We apply new fast-estimation tools to fit mixed effects within a random walk movement model, rapidly inferring among-individual variability in southern elephant seal environment-movement relationships. We found that seals making foraging trips to the sea-ice on or near the Antarctic continental shelf consistently reduced speed and directionality (move persistence) with increasing sea ice coverage and had variable responses to chlorophyll *a* concentration, whereas seals that foraged pelagically reduced move persistence in regions where circumpolar deep water shoaled. Given future climate scenarios, pelagic foragers may encounter more productive habitat but sea-ice foragers may see reduced habitat availability. Our approach is scalable to large telemetry data sets and allows flexible combinations of mixed effects to be evaluated via model selection, thereby illuminating the ecological context of animal movements that underlie habitat use.

## Introduction

Long-term environmental change is influencing southern elephant seal (*Mirounga leonina*) populations, with their trajectories linked to the success of individuals’ foraging migrations (Hindell et al., 2017). These seals migrate long distances from breeding colonies to forage, encountering a range of environmental conditions during many months at sea (Hindell et al., 2017). Foraging strategies vary among seals and are often associated with open ocean or Antarctic continental shelf habitats, with individuals showing fidelity to these over several years (Authier et al., 2012). Quantifying how individuals differently respond to their environment is a challenge due to a paucity of accessible analytical tools that can account for among-individual differences in movement patterns.

Spatial habitat modelling approaches often are used to infer habitat usage and preference from animal movement data (Aarts et al., 2008). Most of these approaches infer preference or selectivity from a combination of observed (presence) and simulated (pseudo-absence) locations (Aarts et al., 2008) but are blind to the ecological mechanisms, such as density dependence (McLoughlin et al., 2010) and individual behaviour (Bestley et al., 2013;Auger-Méthé et al., 2017), underlying those preferences.

Although high individual variation is common in studies of animal movement, models that account for among-individual variability in inferred movement - environment relationships are rare (e.g., McClintock et al., 2013). These random effects or hierarchical models can be computationally demanding, inhibiting realistic analysis of ever-growing animal movement data sets. There is a need for efficient movement modelling approaches, accessible to ecologists, where responses to environmental, physiological and/or social predictors can be inferred using flexible combinations of fixed and random terms (mixed effects) to account for variability among moderate to large numbers (10’s - 100’s) of individuals.

We present a mixed-effects modelling approach for animal movement data that takes advantage of new fast-estimation tools. Our model estimates time-varying movement persistence (autocorrelation in speed and directionality) along animal movement trajectories. We focus here on showing how the approach can be used to infer relationships between animal movement patterns and the environmental features they encounter. The model can be fit rapidly and flexibly with single or multiple random effects, enabling inference across individuals and assessment of the extent to which relationships may differ among them. We apply our approach to infer how southern elephant seals engaging different foraging strategies, ice-bound versus open ocean (pelagic) trips, may respond differently to their environment. This represents a step towards bridging models of animal movement and habitat preference, which in future may converge in a more complete framework.

## Methods

We build our modelling approach in three steps. First, we define a basic model that can be used to estimate changes in move persistence along an animal’s observed trajectory. Second, we expand the model to infer how these changes may be related to environmental variables. Any combination of other extrinsic or intrinsic variables could be modelled, provided they are measured at locations and/or times consistent with the telemetry data. Third, we add random effects to the model to enable inference about how these movement-environment relationships may differ among individual animals.

### Time-varying move persistence

We focus on estimating the persistence (sensu Patlak, 1953) of consecutive pairs of animal relocations (steps) along an entire movement trajectory. Move persistence, which captures autocorrelation in both speed and direction, has been modelled as an average across entire movement trajectories (Jonsen, 2016), indicating whether that trajectory is, on average, uncorrelated (i.e., a simple random walk), correlated (i.e., a correlated random walk), or somewhere in between. Allowing move persistence to vary along a trajectory means it can be used as an index of behaviour (Breed et al., 2012), identifying segments of relatively low or high persistence:

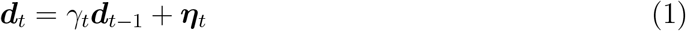

where displacements ***d****_t_* = ***x****_t_* − ***x***_*t*−1_ and ***d***_*t*−1_ = ***x***_*t*−1_ − ***x***_*t*−2_ are the changes in an animal’s location ***x*** at times *t* and *t* − 1. The random variable ***η****_t_* = N(**0**, **Σ**), with variance-covariance matrix **Σ** specifying the magnitude of variability in the 2-dimensional movements. *γ_t_* is the time-varying move persistence between displacements ***d****_t_* and ***d***_*t*−1_. *γ_t_* is continuous-valued between 0 (low move persistence, Appendix S1: Figure S1a,c) and 1 (high move persistence, Appendix S1: Figure S1b,c). To avoid potential parameter identifiability issues between *γ_t_* and **Σ**, we set the covariance term in **Σ** to 0 but this constraint could be relaxed to better account for correlation in movements in the E-W and N-S directions. We assume *γ_t_* follows a simple random walk in logit space:

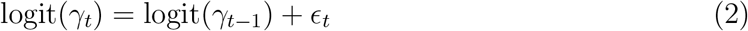

where the random variable *ϵ*_*t*_ = N(0*, σ_γ_*) represents variability in move persistence along an animal’s track.

This process model (Eqn’s 1 and 2) can be fit: 1) to location data with minimal error; 2) to state-space filtered location data; or 3) coupled with an observation model for error-prone data. We focus on the second case with locations occurring at regular time intervals, but this could be relaxed (e.g., Auger-Méthé et al., 2017).

The time-varying move persistence model can be used to objectively identify changes in movement pattern. Here *γ_t_* forms the behavioural index but unlike switching models (e.g., Michelot et al., 2017), these changes occur along a continuum (0 - 1) rather than as switches between discrete states.

### Move persistence in relation to environment

To make inferences about the factors associated with move persistence, we can model *γ_t_* as a linear function of environmental predictors measured at each location or time. With this approach, we replace the random walk on logit(*γ_t_*) (Eqn 2) with a linear regression of covariates on logit(*γ_t_*):

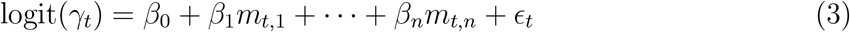

where *β*_0_, *β*_1_,…, *β_n_* are the fixed intercept and regression coefficients, *m_t,_*_1_,…, *m_t,n_* are the predictor variables and *ϵ*_*t*_ = N(0, *σ_γ_*) are the random errors. This model can be fit to a single animal track, or to multiple tracks pooled together. Typically, we wish to make inference across multiple individual tracks and assess the extent to which relationships may differ among individuals.

### Incorporating individual variability

To account for variation among individual responses to environment, we can expand Eqn 3 to a mixed-effects regression of covariates on logit(*γ_t_*), within the behavioural model:

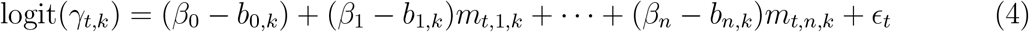

where *k* indexes individual animals, the *β*’s are the fixed intercept and slope terms as in Eqn 3, *b*_0,*k*_ is a random deviation for the intercept of the *k*-th individual, *b*_1,*k*_,…, *b_n,k_* are random deviations for the slopes of the *k*-th individual and *m_t,_*_1,*k*_,…, *m_t,n,k_* are the covariates measured along the *k*-th individual’s track. As in Eqn 3, the random variable *ϵ*_*t*_ are the fixed effects errors. We use an unstructured covariance matrix for the random effects.

### Estimation

In principle, any combination of fixed and random effects can be specified within the movement model described in equations 1 and 4. Here we use TMB to fit the move persistence models (Auger-Méthé et al., 2017). The TMB package allows complex latent variable mixed effects models to be specified in C++ and fit efficiently via maximum likelihood using reverse-mode auto-differentiation and the Laplace approximation (Kristensen et al., 2016). The Laplace approximation avoids the need for high-dimensional integration, which massively speeds calculation of the marginal likelihood. Comparing Bayesian and TMB versions of a location-filtering model, Auger-Méthé et al. (2017) found a 30-fold decrease in computation time for the TMB fit with no loss of accuracy. All code for fitting these models in R is available at https://github.com/ianjonsen.

### Data & analysis

We use Argos telemetry data collected from 24 adult female southern elephant seals. The seals were captured at Iles Kerguelen (49.35° S, 70.22°E) between late January and mid-March in 2009 and 2013-2015, at the end of their annual moult. Animal handling and instrument attachment details can be found elsewhere (McMahon et al., 2008). These data were sourced from the Australian Integrated Marine Observing System (IMOS) deployments at Iles Kerguelen and are publicly available (http://imos.aodn.org.au). The tracks comprise a mixture of sea ice foraging trips on or near the Antarctic continental shelf (12 seals; Appendix S2: Figure S1a) and entirely pelagic foraging trips in sub-Antarctic waters (12 seals; Appendix S2: Figure S1b). Prior to fitting the move persistence models, we used a TMB implementation of a state-space model (Auger-Méthé et al., 2017) to filter the observed locations, accounting for error in the Argos telemetry, and to regularise the filtered locations to a 12-h time interval (see Appendix S2 for details).

We fit the move persistence model (`mpm`; Eqn’s 1 and 2) to the state-space filtered seal tracks. Fitting to filtered tracks accounts for some of the uncertainty inherent in telemetry data but potential effects of residual location uncertainty should be examined post-analysis. To ascertain whether *γ_t_* adequately captures changes in the seals’ movement patterns, we compare the *γ_t_*-based behavioural index to discrete behavioural states estimated from a switching state-space model (Jonsen, 2016) fitted using the bsam R package. Details on how we fit the bsam model are in Appendix S3. We then fit the move persistence mixed effects model (`mpmm`; Eqn’s 1 and 4) to the same state-space filtered seal tracks to infer how the seals’ movement behaviour may be influenced by environmental features encountered during their months-long foraging trips. In both analyses, we fitted separate models to the ice and pelagic foraging trips. For the `mpmm`’s, we specified mixed effects models with random intercept and slopes to account for variability among individual seals. We fit all possible combinations of fixed and random effects and use AIC and likelihood ratios to find the best supported model for each set of tracks.

We examined 3 potential environmental correlates of elephant seal move persistence: sea ice cover (the proportion of time the ocean is covered by ≥ 85% ice; ice), chlorophyll *a* concentration (near-surface summer climatology in mg m^−3^; chl) and the salinity difference between 600 and 200 m depths (based on winter climatology averaged over 1955-2012 in psu; saldiff). These variables are known predictors of elephant seal habitat preference (Hindell et al., 2017) and foraging (McMahon et al. *unpublished data*). Data sources and processing details are provided in Appendix S3. The environmental data values were extracted at each state-space filtered location. As saldiff is only calculated in areas where the bathymetry is deeper than 600 m this covariate is only relevant to the pelagic foragers (Appendix S4: Figure S1). Similarly, ice was excluded from the models fit to seals making pelagic foraging trips as they spent little time in regions with sea-ice cover (Appendix S3: Figure S1; Appendix S4: Figure S1). R code for the model selection is in Appendix S5.

## Results

### Time-varying move persistence (mpm)

The ice-bound seals exhibited similar movement patterns (Fig. 1a), with high move persistence on their outbound migrations and lower move persistence near the Antarctic continent in areas of higher sea-ice coverage. Return migrations to Iles Kerguelen were more variable, with some individuals moving persistently and others meandering, possibly foraging en route. Pelagic foraging seals (Fig. 1b) migrated approximately 2000 km either east or west of Iles Kerguelen in relatively persistent fashion. Less persistent movements occurred at the distal ends of these migrations, although seals travelling to the west of Iles Kerguelen had markedly less persistent and slower movements, suggestive of more intense search and foraging, compared to those travelling to the east (Fig. 1b).

**Figure 1.**
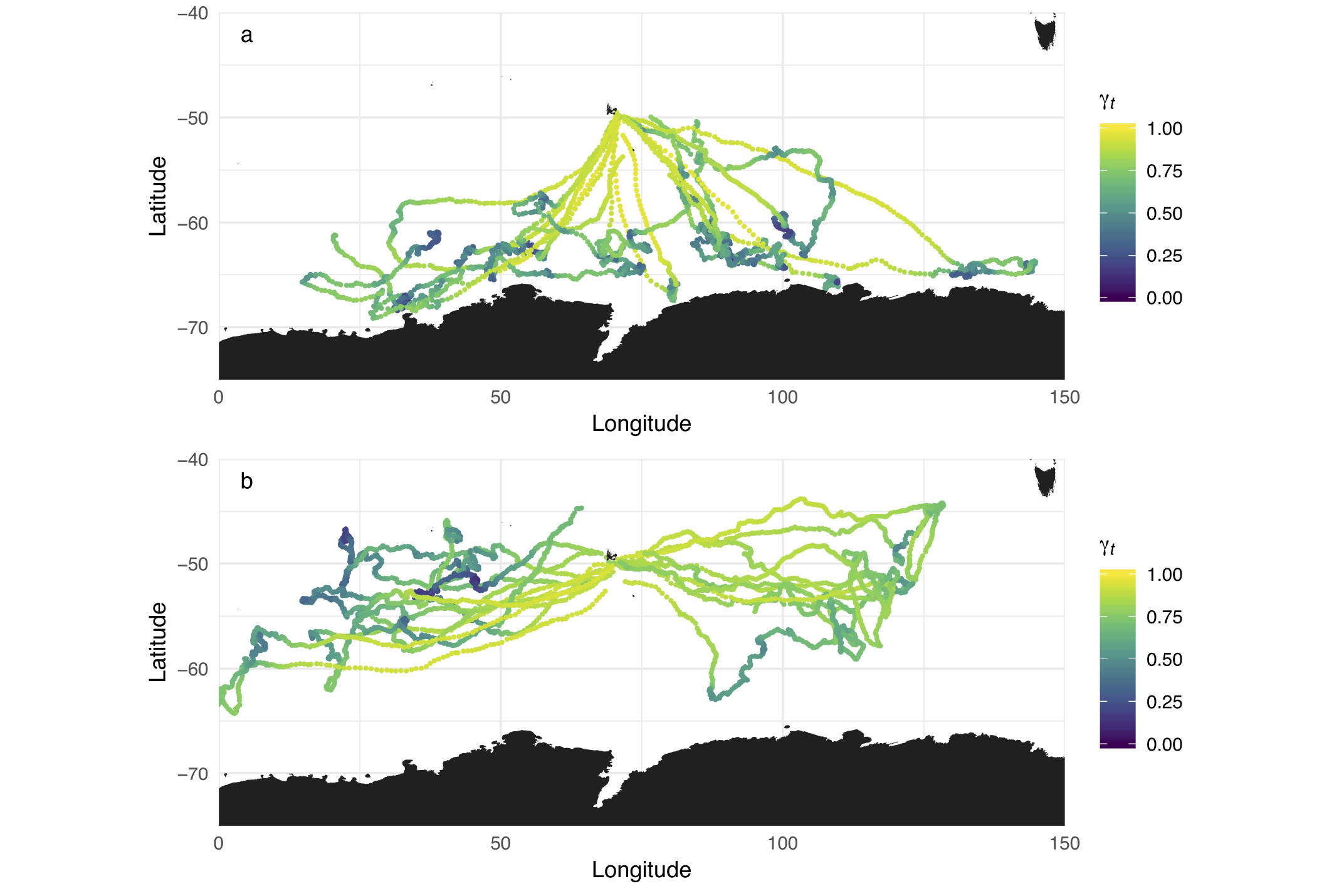
Maps of SSM-filtered southern elephant seal tracks originating from Iles Kerguelen. Ice-bound foraging trips (a) were predominantly directed to locations south of 60°S, whereas pelagic foraging trips (b) are predominantly north of 60°S. Each location is coloured according to its associated move persistence (see *γ_t_* scale bar) estimated from the move persistence model.

The *γ_t_*-derived behavioural index is comparable but not identical to the discrete behavioural states estimated from the `bsam` SSSM (Appendix S3: Figure S1). The *γ_t_* index captured the same changes in movement pattern but the magnitudes of those changes generally were smaller. Fitting the move persistence model, including the SSM filtering step, was almost 500 times faster than fitting the bsam SSSM (Appendix S3: Table S1).

### Individual variability in move persistence - environment relationships (mpmm)

#### Sea-ice strategy

The best supported model for elephant seals foraging in the sea-ice zone included fixed and random coefficients for both the proportion of ice cover and chlorophyll *a* concentration (Table 1). On average, seals had movements that became less persistent or directed as sea-ice cover and chlorophyll *a* concentration increased (Fig. 2a,b). Among individuals, the relationship with ice was consistently negative but the degree to which move persistence declined differed markedly (Fig. 2a), whereas the relationship with chl was highly variable with 4 individuals having strong negative relationships and the rest weak to moderately positive relationships (Fig. 2b; Z-value = −1.04, p = 0.3). Using the fixed effects from the best model, the prediction of *γ_t_* over the spatial domain implies that seal move persistence changes, suggestive of search and foraging behaviours, south of 65°S (south of the black contour line, Fig. 2d).

**Figure 2.**
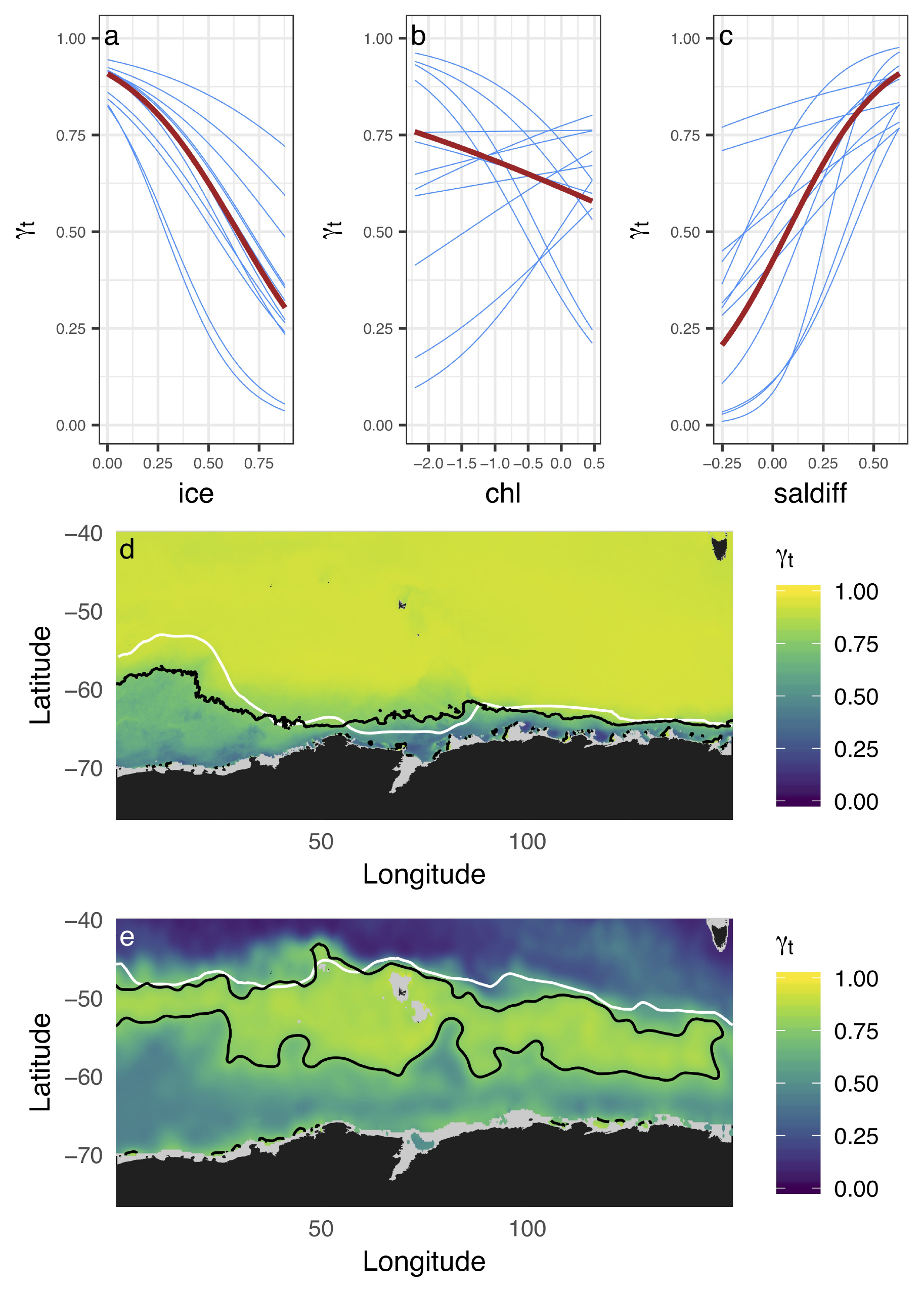
Fixed (red) and random (blue) effects relationships between move persistence *γ_t_* and the proportion of ice cover (a) and chlorophyll *a* concentration (b) for ice foraging seals, and between *γ_t_* and the salinity difference between 600 and 200m (c) for pelagic foraging seals. All three panels display both random intercept and slopes, as per the best ranked models in Table 1. Spatial predictions of *γ_t_* based on the fixed effect coefficients for the best fitting models for ice foraging seals (d) and pelagic foraging seals (e). The *γ_t_* = 0.75 contour (black line) is displayed to aid delineation of predicted high move persistence (*γ_t_* > 0.75; green - yellow) and low move persistence regions (*γ_t_* ≤ 0.75; green - blue). The southern boundaries of the Antarctic Circumpolar Current (d) and the Subantarctic Front (e) are displayed for reference (white lines).

**Table 1:** Model rankings by ∆AIC and likelihood ratios (LR) for the mpmm’s fit to the 24 foraging seals, split by foraging strategy (sea-ice or pelagic). Absolute AIC and deviance values for the best ranked model are displayed on the first row, under the ∆AIC and LR headings. All other ∆AIC and LR values are relative to the best ranked model. Computation time to convergence is also reported. Random effects are included in parentheses in the model formulas.

| Foraging strategy | Model formula | df | $\Delta\text{AIC}$ | LR | Time (s) |
| --- | --- | --- | --- | --- | --- |
| sea-ice | $\sim \text{ice} + \text{chl} + (\text{ice} + \text{chl} \mid \text{id})$ | 12 | -9954.21 | -9978.21 | 4.76 |
| | $\sim \text{ice} + \text{chl} + (\text{chl} \mid \text{id})$ | 9 | 0.78 | 6.78 | 3.61 |
| | $\sim \text{ice} + \text{chl} + (1 \mid \text{id})$ | 7 | 21.06 | 31.06 | 4.17 |
| | $\sim \text{ice} + (1 \mid \text{id})$ | 6 | 21.08 | 33.08 | 2.63 |
| | $\sim \text{ice} + \text{chl} + (\text{ice} \mid \text{id})$ | 9 | 23.59 | 29.59 | 5.76 |
| | $\sim \text{ice} + (\text{ice} \mid \text{id})$ | 8 | 24.14 | 32.14 | 4.55 |
| | $\sim \text{chl} + (\text{chl} \mid \text{id})$ | 8 | 219.74 | 227.74 | 4.09 |
| | $\sim \text{chl} + (1 \mid \text{id})$ | 6 | 245.16 | 257.16 | 3.48 |
| | $\sim 1 + (1 \mid \text{id})$ | 5 | 339.28 | 353.28 | 2.79 |
| pelagic | $\sim \text{saldiff} + (\text{saldiff} \mid \text{id})$ | 8 | -13897.26 | -13913.26 | 3.87 |
| | $\sim \text{saldiff} + \text{chl} + (\text{saldiff} \mid \text{id})$ | 9 | 1.68 | -0.32 | 4.96 |
| | $\sim \text{saldiff} + \text{chl} + (\text{chl} \mid \text{id})$ | 9 | 3.25 | 1.25 | 3.97 |
| | $\sim \text{saldiff} + \text{chl} + (1 \mid \text{id})$ | 7 | 29.81 | 31.81 | 4.04 |
| | $\sim \text{saldiff} + (1 \mid \text{id})$ | 6 | 36.35 | 40.35 | 3.21 |
| | $\sim \text{chl} + (\text{chl} \mid \text{id})$ | 8 | 51.37 | 51.37 | 4.54 |
| | $\sim \text{chl} + (1 \mid \text{id})$ | 6 | 107.41 | 111.41 | 4.19 |
| | $\sim 1 + (1 \mid \text{id})$ | 5 | 129.93 | 135.93 | 2.34 |
| | $\sim \text{saldiff} + \text{chl} + (\text{saldiff} + \text{chl} \mid \text{id})$ | 12 | NA* | NA* | 6.02 |

\*model failed to converge

#### Pelagic strategy

The best supported model for elephant seals foraging in the open ocean included fixed and random coefficients for the salinity difference between 600 and 200 m depths (saldiff, Table 1). On average, seals had movements that became strongly less persistent as the salinity difference decreased (Fig. 2c). Among individuals, this relationship was moderately variable with two individuals exhibiting relatively small changes in move persistence over the full range of saldiff (Fig. 2c). The spatial prediction of *γ_t_* implies that animals generally adopt a movement pattern suggestive of search or forage once beyond the mid-latitudes near Kerguelen Island where saldiff is largest (i.e. south of the black contour line, in oceanic waters, or north in the vicinity of the Subantarctic Front; Fig. 2e).

## Discussion

Southern elephant seals employing specific foraging strategies respond to different environmental factors. Our modelling approach clearly identifies these responses, including strong decreases in move persistence associated with increasing ice coverage (sea-ice foragers) and decreasing salinity difference (pelagic foragers). Move persistence responses were relatively consistent among seals adopting either a sea-ice or a pelagic foraging strategy, but substantial individual variability in foraging location was evident.

Those animals whose forage migrations went towards the Antarctic continent showed low move persistence once in areas of higher sea ice coverage. Some individuals also showed positive responses to elevated chlorophyll *a* concentrations, targeting productive coastal polynya areas (Labrousse et al., 2018); however this was not a consistent response with many others foraging farther offshore in the marginal sea-ice zone where chlorophyll *a* concentrations are lower (Appendix S4: Figure S1). This pattern might be suggestive of density-dependent habitat selection, whereby seals distribute themselves so that foraging success is consistent across habitats of differing value (Morris, 2011).

For the pelagic foraging animals, our results indicated seals moved persistently away from the region in which salty Circumpolar Deep Water was confined to depths (i.e., where the salinity difference was highly positive). The majority then adopted a lower move persistence in areas where the Circumpolar Deep Water shoaled (salinity difference closer to zero, southern areas) with four animals targeting the vicinity of the Subantarctic Front (salinity difference negative) where cold, fresh Antarctic Intermediate Water subducts saline Subantarctic surface waters (northwestern areas, Appendix S4: Figure S1).

Future climate scenarios project stronger westerly winds, leading to intensified ocean overturning circulation (Gao et al., 2018, and references therein). With increased upwelling of nutrient-rich Circumpolar Deep Water, we might expect enhanced near-surface ocean productivity to benefit pelagically foraging southern elephant seals in future. Expectations for sea-ice foraging seals are highly uncertain due to complex physical processes occurring over the Antarctic continental shelf. However, projections of reduced sea-ice extent and duration may lead to reduced availability of foraging and/or resting habitat.

While the ultimate source of observed individual differences in movement - environment relationships is often unclear, three non-exclusive explanations seem likely. First, we often use relatively few predictors and these may indirectly or imperfectly represent the proximate influences to which predators are actually responding (i.e., prey density and/or distribution). This may inflate apparent individual differences in predator movement. Modelling more direct indices of prey, and/or reducing error within covariates by accounting for location uncertainty, may help to reduce apparent variation among individuals.

Second, individual variation is likely a real feature of foraging ecology (Magurran, 1993), where individual quality and personality may confer real differences in foraging behaviour with relatively little difference in fitness (Stamps, 2007). For example, consistent boldness in foraging can generate important ecological trade-offs, effecting increases in growth and/or mortality rates (Stamps, 2007).

Third, the inclusion of multiple random effects raises the possibility of over-fitting, especially when the number of individual tracks is low. Artificial variability, propagating from uncertainty in the locations and/or environmental covariates, could lead to spurious inference of strong individual differences in foraging behaviour. A study design with repeat tagging of the same individuals would help resolve the issue. Ultimately, researchers must take care to address potential sources of error in their data and to use prior knowledge of their study species to guide model selection and interpretation.

Interpreting among-individual variability in movement - environment responses can be aided by considering established ecological theory. For example, density-dependent habitat selection and functional responses to prey availability likely underpin inferred relationships (Mason and Fortin, 2017). Accounting for such effects when fitting and interpreting resource selection functions and habitat preference models can clarify understanding and thereby assist forecasting of species’ distributions (McLoughlin et al., 2010).

### Modelling approach and extensions

Our model is composed of a linear mixed effects regression embedded within a correlated random walk process model for animal movement behaviour. While the linear mixed effects approach allows flexible combinations of fixed and random effects, there is scope for further enhancement. In many cases parametric, linear fixed effects may not adequately capture the complexity of movement - environment relationships and a nonparametric approach using penalised splines may improve inference (Langrock et al., 2017). Given the serial dependence structure of telemetry data, the unstructured covariance matrix we used for the random effects could be replaced with a first-order autoregressive covariance structure (Brooks et al., 2017). Diagnosing lack of fit in latent variable models can be problematic as there is no observed response variable. One-step-ahead prediction residuals provide a useful validation tool and can be estimated when fitting the model (Thygesen et al., 2017). Finally, there is a need to incorporate location uncertainty when sampling environmental covariates from spatially gridded remote-sensing data. This can be done using multiple imputation methods as implemented in momentuHMM R package (McClintock and Michelot, 2018), i.e., random draws of the environmental variables from the uncertainty of the state-space filtered location estimates.

Recent advances in habitat modelling methods (e.g., Avgar et al., 2016) hold promise for capturing the currently missing behavioural context in species’ habitat preferences and space-use. Here we model animal movement as a mixed effects function of environmental variables to gain deeper insight into how individuals and populations may actually use habitat. Our approach does not account for availability/accessibility of habitat in any way but this clearly must be considered when inferring habitat preferences. A reasonable approach for this might be to simulate animal tracks from our movement process model, examining implications of including/excluding environmental covariates. Pseudo-absence tracks can be combined into a habitat accessibility surface to condition spatial prediction of animal behaviour from our process model (e.g., Raymond et al., 2015).

Our results show that TMB allows fast estimation of multiple fixed and random effects in an animal movement process model. Dramatically faster computation times allow analyses of movement - environment relationships in large telemetry data sets (100’s of animals). This computation speed also opens up possibilities for more realistic models of animal movement, where warranted, perhaps by incorporating the third dimension for diving or flying animals and/or high-volume accelerometry data.

The process model used here differs markedly from the state-space model used by Bestley et al. (2013). Bestley et al. (2013) used discrete behavioural state Markov-switching embedded in a correlated random walk process model (Jonsen, 2016). Here, we used time-varying move persistence *γ_t_* as a behavioural index that varied continuously between 0 and 1. This continuous index provides another tool for characterising animal movement patterns and for making inferences about their environmental drivers. In some cases, a continuous index may offer more nuanced insight into variable but unknown behavioural sequences (Breed et al., 2012), whereas discrete states may offer more flexibility in capturing the known structure of animal movement patterns (Michelot et al., 2017).

Telemetry data obtained at the level of individuals poses a challenge to scale up to populations (Morales et al., 2010). Our approach enables multiple fixed (population) and random (individual) effects in movement - environment relationships to be fit simply and quickly. This provides a feasible solution to analysing increasingly large and detailed data sets, and for harnessing individual-to-population level information on animal movement responses to environment.

## Acknowledgements

T Michelot, T Photopoulou, U Thygesen, and S Wotherspoon provided valuable insights that improved the approach and, along with B. Kotler and 4 anonymous reviewers, enhanced the manuscript. This work was supported by: the Climate Impacts on Top Predators (CLIOTOP) Task Team 2016-06 on Animal Movement and Prediction; the Centre for the Synthesis and Analysis of Biodiversity (CESAB); a Macquarie University Co-funded Fellowship to IDJ; an Australian Research Council DECRA grant (DE180100828) to SB; and a CSIRO Julius Career Award and the Villum Foundation to TAP. The Integrated Marine Observing System (IMOS) supported seal fieldwork. IMOS is a national collaborative research infrastructure, supported by the Australian Government and operated by a consortium of institutions as an unincorporated joint venture, with the University of Tasmania as Lead Agent. All tagging procedures approved and executed under University of Tasmania Animal Ethics Committee guidelines.

